# A gene for genetic background in *Zea mays*: fine-mapping *enhancer of teosinte branched1.2* (*etb1.2*) to a YABBY class transcription factor

**DOI:** 10.1101/070201

**Authors:** Chin Jian Yang, Lisa E. Kursel, Anthony J. Studer, Madelaine E. Bartlett, Clinton J. Whipple, John F. Doebley

**Author notes:** Present address: Molecular & Cellular Biology Graduate Program, University of Washington, Seattle, WA 98195. Present address: Department of Crop Sciences, University of Illinois Urbana-Champaign, Urbana, IL 61801. Present address: Department of Biology, University of Massachusetts Amherst, Amherst, MA 01003. Corresponding author: Laboratory of Genetics, University of Wisconsin-Madison, Madison, WI 53706.

## Abstract

The effects of an allelic substitution at a gene often depend critically on genetic background, the genotype at other genes in the genome. During the domestication of maize from its wild ancestor (teosinte), an allelic substitution at *teosinte branched* (*tb1*) caused changes in both plant and ear architecture. The effects of *tb1* on phenotype were shown to depend on multiple background loci including one called enhancer of *tb1.2 (etb1.2).* We mapped *etb1.2* to a YABBY class transcription factor (*ZmYAB2.1*) and showed that the maize alleles of *ZmYAB2.1* are either expressed at a lower level than teosinte alleles or disrupted by insertions in the sequences. *tb1* and *etb1.2* interact epistatically to control the length of internodes within the maize ear which affects how densely the kernels are packed on the ear. The interaction effect is also observed at the level of gene expression with *tb1* acting as a repressor of *ZmYAB2.1* expression. Curiously, *ZmYAB2.1* was previously identified as a candidate gene for another domestication trait in maize, non-shattering ears. Consistent with this proposed role,*ZmYAB2.1* is expressed in a narrow band of cells in immature ears that appears to represent a vestigial abscission (shattering) zone. Expression in this band of cells may also underlie the effect on internode elongation. The identification of *ZmYAB2.1* as a background factor interacting with *tb1* is a first step toward a gene-level understanding of how *tb1* and the background within which it works evolved in concert during maize domestication.

---

The effects of an allelic substitution at a gene can vary widely depending on the genetic background in which its effects are assayed (Chandler *et al.* 2013). In extreme cases, the effect of an allele may even exhibit a change in directionality across different genetic backgrounds (Wade *et al*. 2001). Often, genetic background is considered a “nuisance factor” and thus something that needs to be controlled by the usage of inbred lines in experiments involving model organisms such as maize, *Arabidcrpsis* and mice because the effect of an allele can change under different genetic backgrounds (Wade 2002; Chandler *et al.* 2013). However, in evolutionary biology, a knowledge of the extent and dynamics of genetic background is essential for a complete understanding of the evolutionary process.

The *teosinte branched* (*tb1*) gene of maize (*Zea mays ssp. mays*) has been identified as one of the genes involved in the domestication of maize from its wild progenitor, teosinte (*Zea mays ssp. parviglumis*). *tb1* acts pleiotropically to regulate both the architecture of the plant overall and the structure of the female inflorescence or ear (Doebley *et al.* 1997). Plants carrying the maize allele of *tb1* have fewer, shorter branches than plants carrying teosinte alleles. The ears of plants with the maize allele have short internodes such that the kernels are more tightly packed together as compared to plants with teosinte alleles. Substitution of a teosinte allele for a maize allele of *tb1* can also partially transform the ear (female) into a tassel (male).

The effects of the maize vs. teosinte alleles of *tb1* on both plant and ear traits vary with genetic background, and epistatic interactions between *tb1* and other loci that contribute to the background effects have been described (Doebley *et al*. 1995; Studer and Doebley 2011). Among these other loci is a quantitative trait locus (QTL) called *enhancer of tb1.2 (etb1.2*). The principal effect of *etb1.2* is on the length of internodes within the ear, although it also affects the sex of the ear itself, whether partially converted into a tassel by the teosinte allele. Interestingly, *etb1.2* is located only 6 cM upstream of *tbl.1*

Here, we report the fine-mapping of *etb1.2* to a YABBY family transcriptional regulator called *ZmYAB2.1.* Expression assays show that some maize alleles of *etb1.2* are expressed at a lower level than teosinte alleles while other maize alleles are loss-of-function with disrupted coding sequences. There is an epistatic interaction between *etb1.2* and *tb1* such that the effects of the maize vs. teosinte allele of *etb1.2* are much stronger when *tb1* is fixed for the teosinte allele. This interaction effect is seen both at the level of phenotype and *ZmYAB2.1* expression. *tb1* functions as an upstream regulator (repressor) of *etb1.2.* Surprisingly, *ZmYAB2.1* has been previously implicated as a gene controlling another domestication trait - non-shattering inflorescences (Lin *et al.* 2012). Consistent with an effect of *ZmYAB2.1* on shattering, we show that *ZmYAB2.1* is expressed in a narrow band of cells that likely represents a vestigial shattering (abscission) zone. Expression in this band of cells may also explain the effect on ear internode length. This report provides the first gene-level view of how genetic background influences the effects of *tb1*.

## Materials and Methods

### Finemapping of etb1.2

We constructed 45 recombinant chromosome nearly isogenic lines (RC-NILs) carrying different teosinte (*Zea mays ssp. parviglumis;* Iltis and Cochrane collection 81) introgressions within the interval of *etb1.2* QTL (Studer and Doebley 2011) for fine-mapping. We obtained these RC-NILs by first crossing two nearly isogenic recombinant inbred lines (NI-RILs) with the teosinte allele at *tb1* and maize W22 background but different alleles (maize versus teosinte) at *etb1.2* (Supporting information, Figure S1). We then screened 1425 plants from the resulting F2 population for recombinants in the *etb1.2* interval using two flanking markers: *umc2047* and *umc1411.* Plants with recombinant chromosomes were then selfed to produce the 45 RC-NILs. We further narrowed the recombination breakpoints of each RC-NIL using genotype-by-sequencing (GBS) markers (Elshire *et al.* 2011), Sanger sequencing-based SNP markers, and insertion/deletion-based markers (Supporting information, Table S1).

We planted all 45 RC-NILs and the two NI-RIL parents using a randomized complete block design at the University of Wisconsin West Madison Agricultural Research Station in Summer 2012. These lines were planted in 6 blocks with 68 plots in each block, where 24 RC-NILs had one seed-lot planted and 21 other RC-NILs had two seed-lots planted. Each plot was 3.6m long and 0.9m wide, and was sowed with 12 seeds. For each plot, we harvested the top ear from 5 plants for scoring the ear internode length (length of 4 to 10 consecutive kernels divided by the number of kernels in millimeters). This trait was also known as average length of the cupules in the ear (CUPL) or 10-kernel length (10KL) in previous publications (Doebley *et al*. 1995; Studer and Doebley 2011).

We computed the least square means (LSMs) of ear internode length to be used in fine-mapping. These LSMs were calculated using the MIXED procedures in SAS (Littell *et al.* 1998), and the model is shown as following:

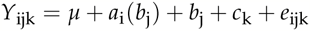

where Y_ijk_ is the ear internode length for an introgression line, *μ*, is the overall mean, a_i_ is the fixed seed-lot effect, b_j_ is the fixed line effect, c_k_ is the random block effect, and e_ijk_ is the experimental error.

To fine-map the interval within which the *etb1.2* QTL is located, we looked for correspondence between the introgressed teosinte chromosomal segments and LSMs of the 45 RC-NILs. We first matched the introgressed segments of the 45 RC-NILs to their corresponding LSMs. Next, we sorted the 45 RC-NILs by their LSMs and identified two distinct classes of LSMs separated by non-overlapping error bars. Under the assumption that there is a single gene contributing to the QTL, one expects a clean segregation of the lines into two groups based on their LSMs such that there is a small chromosomal interval for which all lines in one group have teosinte genotype and all lines in the other group have maize genotype. The interval thus identified is the causative interval that contains the QTL. Genotype and phenotype data are available from the Dryad Digital Repository: (http://dx.doi.org/XXX).

### Gene Characterization

Fine-mapping identified *ZmYAB2.1* as the only gene within the refined QTL interval. We obtained the genomic sequences of *ZmYAB2.1* for B73, W22 and our teosinte parent by Sanger sequencing. Resequencing the B73 allele was necessary because the available published sequences were incomplete. We designed primers for sequencing *ZmYAB2.1* based on GRMZM2G085873 gene model in the maize reference genome AGPv2 (http://www.maizegdb.org) and an improved *ZmYAB2.1* sequence assembly that was published as *ZmSh1* (Lin *et al.* 2012). A complete list of primers used for sequencing can be found in Table S2 (Supporting information). A B73 BAC clone (ZMMBBc0564C07) from Arizona Genome Institute was also used for facilitating the sequencing work. We performed all sequence alignment using Sequencher version 5.1 (Gene Codes Corporation). All our sequences have been deposited in GenBank (Accession Numbers: KX687024 - KX687134).

We performed phylogenetic analysis of the YABBY gene family across diverse plant species using 107 published YABBY peptide sequences. Of these 107 YABBY peptide sequences, 101 were taken from Yamada *et al.* (2011) and 6 other were taken from UniProt (The UniProt Consortium 2015) and Lin *et al.* (2012). We aligned all 107 YABBY peptide sequences using Clustal X2.1 (Larkin *et al.* 2007) and Se-Al v2.0a11 (Rambaut 2002). Conserved positions of the 107 YABBY peptide sequences were extracted using Gblocks 0.91b (Castresana 2000). We performed Bayesian phylogenetic analysis of the extracted YABBY peptide sequences using MrBayes v3.1.2 on XSEDE provided by the CIPRES Science Gateway (Miller *et al.* 2010). Parameters used in this phylogenetic analysis were kept the same as the parameters originally used by Yamada *et al.* (2011). The model for among-site rate variation (rates=) was set to invariant + gamma and rate matrix for amino acids (aamodelpr=) was set to Jones (Jones *et al.* 1992). Trees were sampled at every 1,000 generations (sampfreq=) for 5,000,000 number of generations (ngen=) with the first 25% discarded as burn-in. We then verified the convergence of the Markov Chain Monte Carlo (mcmc) runs using Tracer v1.60 (Rambaut *et al.* 2014).

### Population Genetics

We investigated for evidence of selection during domestication around *ZmYAB2.1* using Hudson-Kreitman-Aguade (HKA) test (Hudson *et al.* 1987), Tajima’s D (Tajima 1989), and coalescent simulation (Innan and Kim 2004). We also calculated the ratio of nucleotide diversity (*π*) in maize as compared to teosinte and the same ratio for nucleotide polymorphism (*θ*). To perform these analyses, we first sequenced the 5’ end of *ZmYAB2.1* (includes 5’ UTR and Exon 1) and two regions upstream of *ZmYAB2.1* from 19 maize landrace accessions, 16 teosinte accessions and 1 *Zea diploperennis* accession (outgroup for HKA test) (Supporting information, Table S3. These three regions were chosen to cover the fine-mapped *etb1.2* interval. We also included sequences from six previously identified neutral genes (AY104395, AY106816, AY107192, AY107248, AY111546, AY111711) (Zhao *et al.* 2011) as control genes for the HKA test. All sequence alignments were done using Sequencher version 5.1 (Gene Codes Corporation). We analyzed nucleotide diversity (*π*), nucleotide polymorphism (*θ*), HKA test (Hudson *et al.* 1987) and Tajima’s D (Tajima 1989) using DnaSP v5.10.01 (Librado and Rozas 2009). Percent loss in diversity was calculated as 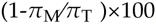. We performed coalescent simulation using software from Innan and Kim (2004) with the same parameters used by Zhao *et al.* (2011).

We investigated the distribution of four null *ZmYAB2.1* alleles in 69 *Zea mays* ssp. *mexicana* accessions, 73 *Zea mays* ssp. *parviglumis* accessions, 127 *Zea mays* ssp. *mays* accessions (landrace), 18 teosinte inbred lines, and 58 maize inbred lines (Supporting information, Table S4. The four null *ZmYAB2.1* alleles were discovered during our diversity sequencing and expression analyses on *ZmYAB2.1* in multiple maize inbred lines. We named these null alleles after the maize inbred line in which they were initially identified and the corresponding mutation, namely B73_Spm, W22_Spm, Mo17_82bp, and Ki3_2bp. We screened for the B73_Spm and W22_Spm alleles using PCR with a three primers mix (Supporting information, Table S5) followed by agarose gel electrophoresis. For each null allele screen, we designed two primers to flank the insertion and a third primer that sits within the insertion. We scored the PCR products based on size differences on a 4% agarose gel. We screened for the Mo17_82bp and Ki3_2bp alleles using PCR with fluorescent primers (Supporting information, Table S5) followed by assaying the PCR products in ABI 3730 DNA Analyzer (Applied Biosystems). We then analyzed the results using Gene Marker version 1.7 (Softgenetics, State College, PA). To sum up all the null alleles, we plotted the geographical distributions of all assayed alleles on map using Google Map Engine Lite (https://mapsengine.google.com/map/) (Supporting information, Figure S2).

### Gene Expression

#### Quantitative RT-PCR

We performed gene expression assays for *ZmYAB2.1* using RNA extracted from immature ears between 10 – 30 mm (roughly 60 days post planting) using the standard Trizol (Invitrogen) protocol. We evaluated the isolated RNA quality using agarose gels made from 3-(N-morpholino)propanesulfonic acid (MOPS) buffer and formaldehyde. Following that, we treated each RNA sample with DNase prior to cDNA synthesis using standard oligo dT primers and Superscript III reverse transcriptase (Invitrogen). We also verified the cDNA quality and absence of DNA contamination by PCR (Taq Core Kit, Qiagen) by amplifying across two exons using flanking primers in GRMZM2G126010 (actin) (Supporting information, Table S5).

We performed quantitative RT-PCR (qPCR) analysis of *ZmYAB2.1* in the two parental NI-RILs (T1L_C07, T1L_D17), carrying either the teosinte introgression or maize (W22) allele at *etb1.2.* Immature ears for these two NI-RILs were harvested from plants grown at the University of Wisconsin West Madison Agricultural Research Station (UW-WMARS) in summer 2013. We specifically designed the PCR primers to span across two or more exons as an insurance against DNA contamination during qPCR. We used two sets of primers to amplify 220bp fragment from *ZmYAB2.1* and 201bp fragment from GRMZM2G126010 (actin gene) respectively (Supporting information, Table S5). We normalized *ZmYAB2.1* expression level based on GRMZM2G126010 (actin gene). All qPCR reactions were done using Power SYBR Green PCR Master Mix (Applied Biosystems) in ABI Prism 7000 Sequence Detection System (Applied Biosystems). The following steps were used in the qPCR reactions: activation at 95°C for 10 minutes, denaturation at 95°C for 15 seconds, annealing and extension at 60°C for 1 minute, and repeated for 40 cycles. A total of 20 biological replicates and 3 technical replicates were used for each NI-RIL in this expression analysis.

Similarly, we also performed qPCR analysis of *ZmYAB2.1* with two RC-NILs: IN1207 carries the teosinte allele of *ZmYAB2.1,* and IN0949 carries a recombinant allele with maize sequence in the 5’ regulatory region and teosinte sequence in the coding region of *ZmYAB2.1* (Supporting information, Figure S3). Immature ears for these two RC-NILs were harvested from plants grown at the University of Wisconsin Walnut Street Greenhouse (UW-WSGH) in winter 2013. We used the same primer sets, reagent, instrument and replicate setup as described in previous paragraph.

#### Allele-Specific Expression Assays

We also investigated the difference in the mRNA abundance of *ZmYAB2.1* between maize and teosinte alleles using an allele specific expression assay. This assays compares mRNA abundance of the two alleles when heterozygous in a plant. Maize *ZmYAB2.1* allele was represented by two maize inbreds (A619 and Oh43) and two maize landraces inbreds (MR12 and MR19) while teosinte *ZmYAB2.1* allele was represented by two teosinte inbreds (TIL01 and TIL11) (Supporting information, Table S4). These lines were chosen because all of them have functional alleles encoding a full length protein (Supporting information, Table S4). RNA was extracted from immature inflorescences at the tip of the lateral branch from the F_1_ hybrid plants grown at the UW-WSGH in Winter 2014.

For each F1 RNA sample, cDNA was synthesized using the method described above. In contrast to qPCR, the allele specific expression analysis requires small insertion/deletion (indel) that distinguishes the two alleles being compared. As a consequence, we designed a set of primers (Supporting information, Table S5) flanking an indel for amplifying a small fragment within the 5’ UTR of *ZmYAB2.1* (Supporting information, Table S6). To perform the allele specific expression analysis, we ran a PCR of the 5’ UTR fragment with fluorescent primers on the cDNA, and assayed the PCR products in ABI 3730 DNA Analyzer (Applied Biosystems). We obtained the areas under the peaks corresponding to maize and teosinte alleles using Gene Marker version 1.7 (Softgenetics, State College, PA). We calculated the relative expression of the alleles by taking the ratio of areas under the peaks for teosinte allele over maize allele. For each F_1_ hybrid, there were between three to eight biological replicates and three technical replicates. Of the 8 possible teosinte x maize F_1_s, only six had different allele sizes for the indel.

Since different alleles may have different efficiencies for PCR amplification, we needed to perform correction on the allele specific expression assay using DNA controls. We used DNA isolated from the F_1_ plants as a control with a known ratio of 50:50 for the teosinte and maize alleles. DNA was isolated from leaf tissue of each plant using standard CTAB extraction protocol (Rogers and Bendich 1985). We included these DNA controls in each technical replicate of the assay. We identified any PCR amplification bias by taking the ratio of areas under peaks for maize allele over teosinte allele, which is also known as the correction factor. We then multiplied the relative expression by the correction factor to obtain the final corrected relative expression of teosinte allele over maize allele. For example, if the ratio of teosinte expression level over maize expression level is 2.0, and the correction factor is 1.5, then the final corrected relative expression would be 3.0 to account of the PCR amplification bias towards maize allele. Gene expression data are available from the Dryad Digital Repository: (http://dx.doi.org/XXX).

### Interactions with tb1

In order to investigate any interaction between *ZmYAB2.1* and *tb1* in the regulation of ear internode length, we selected 20 NI-RILs for phenotypic and expression analyses (Supporting information, Figure S4. For the phenotypic analysis, we planted these 20 NI-RILs using a randomized complete block design at the University of Wisconsin West Madison Agricultural Research Station in Summer 2014. These lines were planted in 6 blocks with 20 plots in each block. Each plot was 3.0 m long and 0.9 m wide and sown with 10 kernels. For each plot, we harvested the uppermost ear from 5 plants for scoring the ear internode length (length in millimeters of 4 to 10 consecutive kernels divided by the number of kernels). We obtained the LSMs for the ear internode length using similar model described above. For the expression analysis, we quantified both *ZmYAB2.1* and *tb1* expression levels in four immature ears from each of these 20 NI-RILs. The protocol for the expression analysis was the same as described above.

We tested for three different interactions between *ZmYAB2.1* and *tb1* genotypes on *ZmYAB2.1* expression, *tb1* expression, and ear internode length. We tested for these interactions using the following linear model implemented in R (R Core Team 2016):

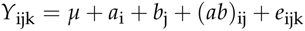

where *μ* is the overall mean, a_i_ is *ZmYAB2.1* genotype main effect, b_j_ is *tb1* genotype main effect, (ab)_ij_ is interaction effect between ZmYAB2.1 and *tb1,* and e_ijk_ is the experimental error. Three different datasets (*ZmYAB2.1* expression, *tb1* expression and ear internode length) were used for Y_ijk_. Phenotype and gene expression data are available from the Dryad Digital Repository: (http://dx.doi.org/XXX).

### in situ Hybridization

Maize ears (20 – 30 mm) and teosinte ears (10 – 20 mm) were collected and fixed in 4% para-formaldehyde, then embedded in paraffin, as described by Jackson (1991). Two digoxygenin-UTP labeled RNA antisense probes were created using MEGAScript T7 kit (Ambion). One probe to the 3’ UTR was amplified with YABBY3’UTRF1 (CCTATGAAAGTAGCAACAAG) and YABBY3’UTRR1 (ATACACGACACTCA-CAAACTA). Another probe containing coding sequence from exons 2 and 3 downstream of an Enhancer/Suppressor Mutator insertion was originally amplified from cDNA with the primers YABBYExon2.3F1 (CGTCCAGAATCACTACTCAC) and YABBYExon2.3R1 (GTGCAGCATATGATCCAGAT). A third probe to the 5’UTR was amplified with YABBY 5’UTRF1 (CTGGATG-GCCCTCTGATTAG) and YABBY5’UTRR1 (ATCTGGGCCGCC-GACGACAT) The 3’UTR probe was mixed 1:1 with either the exon 2/3 probe or the 5’ UTR probe for hybridizations in Figure 4A-D, while only the 3,’UTR probe was used in Figure 4E-F and Figure S6 (Supporting information). Sectioning, hybridization, washing and detections was performed as previously described (Jackson 1991; Jackson *et al.* 1994) except hybridization and washing temperature was raised from the standard 55°C to 65°C for the 3’UTR probe.

## Results

### Fine-mapping of etb1.2

We successfully narrowed *etb1.2* down to a 68 kb interval (AGPv2, chr1: 260,383,398 – 260,448,142) containing exon 1 and 5’ regulatory region of the maize filtered-gene-set member GRMZM2G085873. Among the 45 RC-NILS and two parental NI-RILs, we observed a segregation of the lines into two distinct classes for the least-square means (LSMs) of ear internode length (Figure 1). Examination of the genotypes of these two classes reveals that the class with short internodes carries maize genotype between markers SBM03 and SBM04, and the class with long internodes carries teosinte genotype between these markers (Figure 1). Interestingly, while this interval includes exon 1 of GRMZM2G085873, the only coding sequence difference between the W22 and teosinte alleles is a single synonymous substitution in exon 1 which seems unlikely to confer differences in ear internode length between maize and teosinte. Based on these observations, we hypothesize that a *cis*-regulatory change upstream of GRMZM2G085873 is the likely causal polymorphism underlying *etb1.2.*

**Figure 1.**
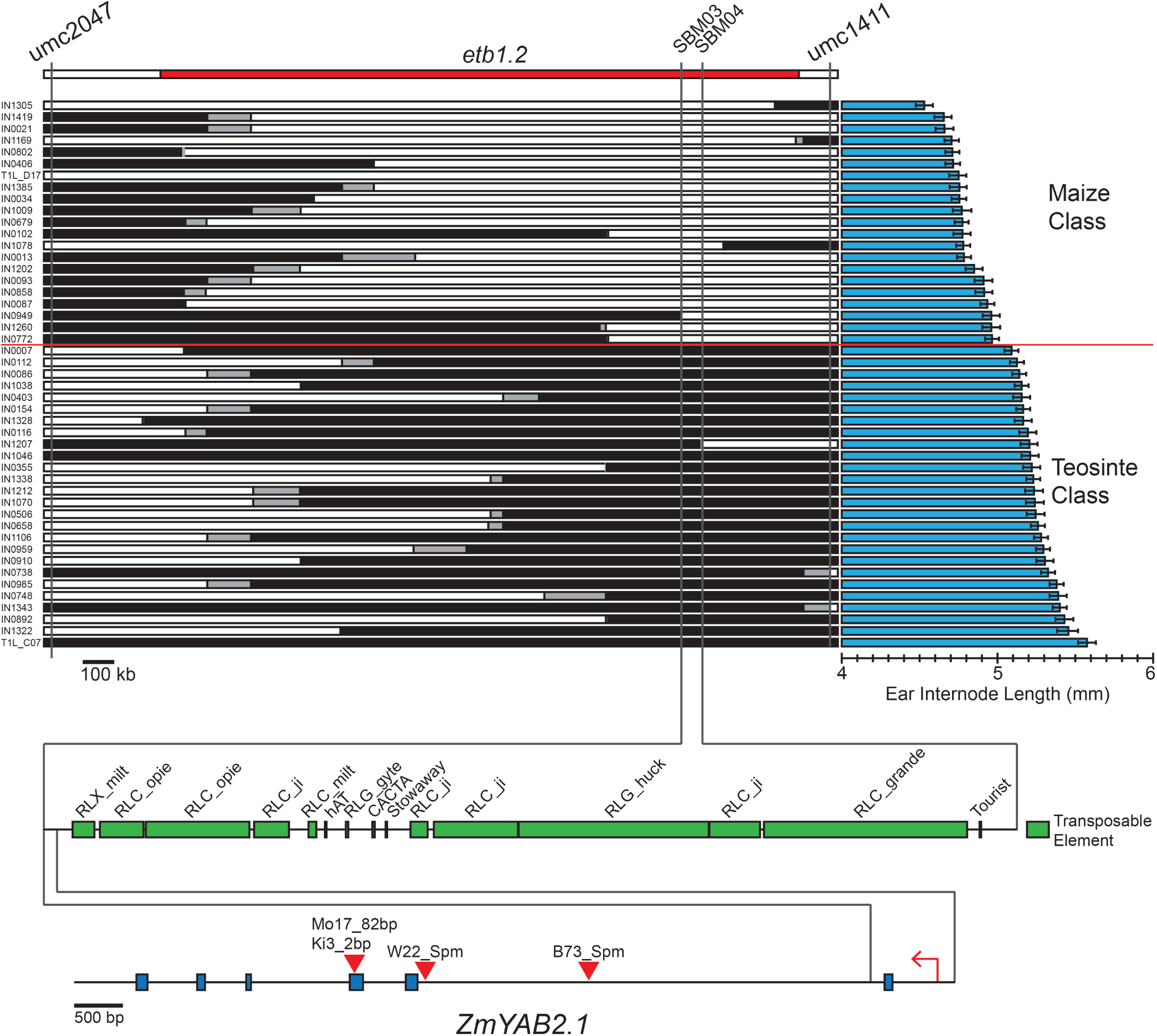
Fine-mapping of *etb1.2* on chromosome 1L. Genotypes of 45 RC-NILs and two NI-RILs carrying different teosinte introgression within *etb1.2* are shown on the left. White bars represent maize genotype, black bars represent teosinte genotype, and grey bars represent unknown genotype. Least squares means and standard errors of ear internode lengths for each line are shown on the right. Markers SBM03 and SBM04 flank the narrowed *etb1.2* interval, which is expanded at the bottom to show a close-up view of the region. Multiple insertion alleles (B73_Spm, W22_Spm, Mo17_82bp, Ki3_2bp) are shown in red triangles.

### Gene Characterization

Examination of the annotation for GRMZM2G085873 revealed that it encodes a protein with homology to the YABBY transcription factor family, first identified in *Arabidopsis.* Based on the maize B73 AGPv2 reference genome, GRMZM2G085873 (260,364,371 – 260,367,349) possesses five exons; however, the *Arabidopsis* YABBY homolog contains six exons. Further examination of the B73 AGPv2 reference sequence shows that the assembly has a contig in the wrong orientation (Supporting information, Figure S6). We re-sequenced this region in B73, and our sequencing results indicate that GRMZM2G085873 actually has 6 exons in B73 like its *Arabidopsis* counterpart (Supporting information, Figure S7). Our sequence data also show that W22 and our teosinte allele possess 6 exons. In our complete alignment of 10,677 bp, the B73 sequence is 10,046 bp, the W22 sequence is 8,993 bp, and teosinte sequence is the 8,362 bp. A BLAST search of the complete sequence using RepeatMasker (Smit *et al.* 2015) on RepBase (Jurka *et al.* 2005) identified a defective Enhancer/Suppressor Mutator element, also known as En/Spm-I or dSpm (Peterson 1953; McClintock 1954), inserted between the first and second exons in B73. We also found a similar, but not the same, insertion in W22 at a different position between the first and second exons (Supporting information, Figure S7).

Phylogenetic analysis revealed that GRMZM2G085873 belongs to the YAB2 clade of the five known clades in the YABBY gene family (Supporting information, Figure S8) (Siegfried *et al*. 1999; Bowman 2000; Yamada *et al*. 2011). Given this result, we designate GRMZM2G085873 as *ZmYAB2.1.* YABBY genes encode transcription factors, which consist of a C_2_C_2_ zinc finger domain that binds DNA and a helix-loop-helix YABBY domain that recognizes DNA binding sites (Bowman 2000). YABBY transcription factors are mainly expressed in the abaxial domain of lateral organs and contribute to adaxial-abaxial polarity (Siegfried *et al.* 1999), although in maize the polarity is switched and they are expressed adaxially (Juarez *et al.* 2004). Vegetatively expressed-YABBY genes (FIL, *YAB2, YAB3,* and *YAB5*) were initially thought to directly specify leaf adaxial-abaxial polarity. However, recent analysis indicates that YABBY genes play a central role in diverse leaf growth and patterning including blade expansion, inhibition of meristematic genes, and signaling to the adjacent meristem (Goldshmidt *et al.* 2008; Sarojam *et al.* 2010).

*ZmYAB2.1* (also called *ZmShl-l*) was previously reported as the ortholog of *Shattering1* of sorghum, a key gene controlling the difference between shattering and non-shattering inflorescences in wild versus domesticated sorghum (Lin *et al.* 2012). In maize-teosinte QTL studies, a QTL for inflorescence shattering mapped to *ZmYAB2.1*, making it a strong candidate for the gene controlling the difference in shattering between teosinte and maize as well (Lin *et al.* 2012). In teosinte, shattering occurs via abscission layers that form at the joints (nodes) between successive internodes in the ear. This prior report and our results suggest that *ZmYAB2.1* may play a dual role of regulating both internode elongation and abscission layer formation for inflorescence shattering.

### Population Genetics

Based on multiple tests for departure from neutral evolution, we identified some evidence for selection near the 5’ end of *ZmYAB2.1,* although the evidence is fairly weak. We did not find a significance departure from neutrality with the Tajima’s D test nor a test based on coalescent simulations in any of the three regions tested; however, we did find a significant departure from neutrality at the 5’ end of *ZmYAB2.1* with the HKA test (Table 1). The 5’ region also shows highest loss in nucleotide diversity in maize as compared to teosinte. Failure of Tajima’s D to reject neutrality is not surprising given that it has been previously shown that Tajima’s D has low power for detecting selection in maize (Tenaillon *et al.* 2004). Similarly, the coalescent simulation test has also been shown to be conservative (Zhao *et al.* 2011). Nonetheless, with some evidence for selection at the 5’ end of *ZmYAB2.1,* the causative site for the difference between maize and teosinte is likely to be close to *ZmYAB2.1* and within the 68 kb interval defined by the fine-mapping.

**Table 1.**
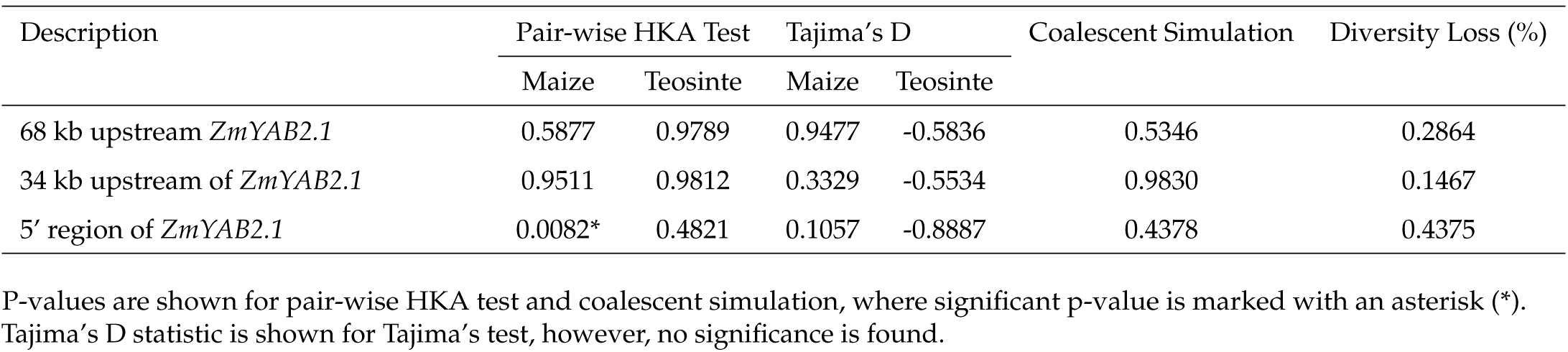
Results for 3 different selection tests on 3 different regions upstream of *ZmYAB2.1*.

We identified four different mutant alleles of *ZmYAB2.1* while sequencing several maize inbred lines to clarify the gene structure. The first two mutant alleles, B73_Spm and W22_Spm, were initially discovered from sequencing of *ZmYAB2.1* in B73 and W22. As described in previous section, both B73 and W22 carry unique dSpm insertions in between the first and second exon. Simple expression analysis via reverse transcription-PCR (RT–PCR) revealed that the *ZmYAB2.1* transcript in B73 and W22 are truncated, including only the first exon (data not shown). We hypothesized that the dSpm insertions are responsible for the truncation of the *ZmYAB2.1* transcripts. Two other mutant alleles, Mo17_82bp and Ki3_2bp, were discovered from sequencing transcripts of *ZmYAB2.1* in Mo17 and Ki3. As the names suggest, Mo17 carries an 82 bp out-of-frame insertion in the third exon of *ZmYAB2.1,* while Ki3 carries a 2 bp out-of-frame insertion in the third exon of *ZmYAB2.1.* All four mutant alleles likely result in functional knockout of *ZmYAB2.1*.

Knowing that maize has multiple mutant alleles of *ZmYAB2.1,* we surveyed the distribution of these mutant alleles in diverse maize and teosinte populations. We found that these four mutant alleles are relatively rare (0 - 2%) in teosinte populations, while three of the mutant alleles are widespread (4 - 22%) in maize populations (Table 2; Supporting information, Table S4). The frequencies and geographic distributions of the mutant allele frequencies suggest that B73_Spm and W22_Spm alleles, which are broadly distributed, arose early during the diversification of maize, while Mo17_82bp and Ki3_2bp alleles arose much later and are geographically confined (Supporting information, Figure S2). B73_Spm alleles are concentrated around South America, W22_Spm alleles are prevalent around Central America, while Mo17 alleles are restricted to North America and the Ki3_2bp allele is known only from that single inbred line.

**Table 2.**
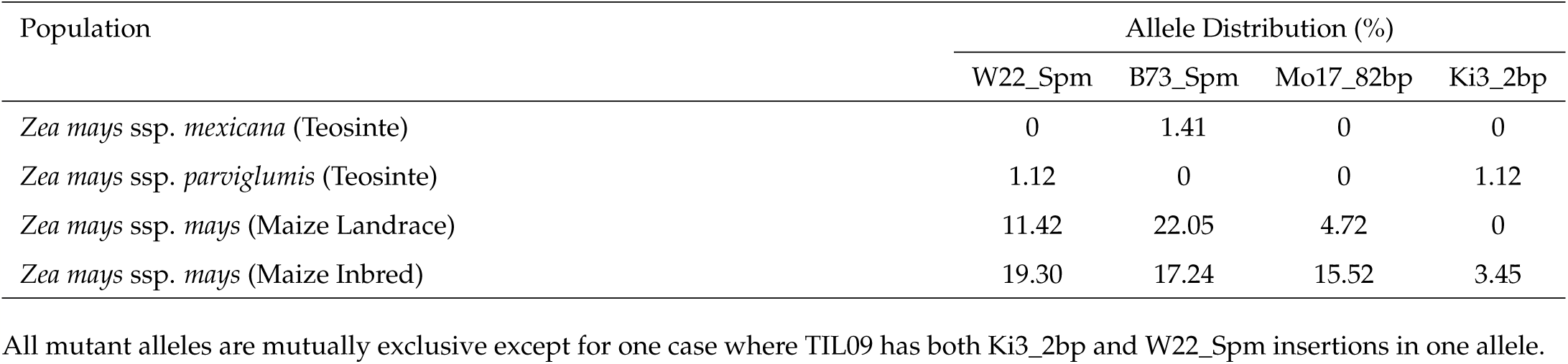
Allele distribution of four different mutant alleles of *ZmYAB2.1* across diverse maize and teosinte populations.

### Gene Expression

Given that we earlier hypothesized that a *cis*-regulatory change at *ZmYAB2.1* is a causal polymorphism underlying *etb1.2,* we first analyzed the expression of the *ZmYAB2.1* maize and teosinte alleles in the two parental lines, T1L_C07 (teosinte allele) and T1L_D17 (W22 maize allele). We found that the *ZmYAB2.1* W22 maize allele (T1L_D17) is not expressed, while the teosinte allele (T1L_C07) is expressed at a relatively high level (Figure 2A). This was the expected result given that the W22 allele has the dSpm insertion which should block the synthesis of a stable transcript. We excluded the possibility of primer mismatch in the W22 maize allele since the primers sequences match perfectly to the W22 maize and teosinte sequences.

**Figure 2.**
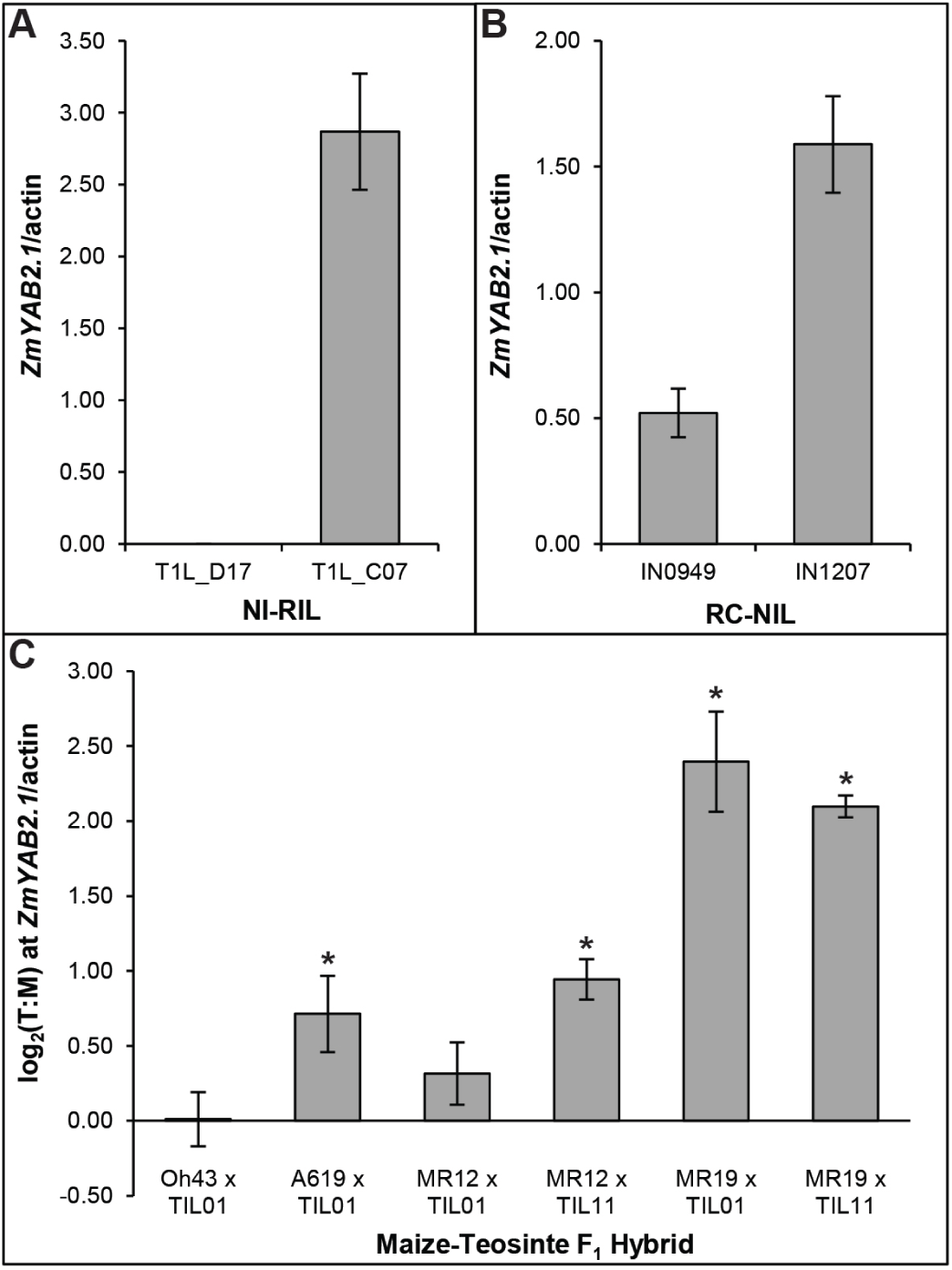
Expression levels of *ZmYAB2.1* in various back-grounds. Teosinte *ZmYAB2.1* allele (T1L_C07) is expressed at a relatively high level while maize *ZmYAB2.1* allele (T1L_D17) is not expressed [A]. Recombinant *ZmYAB2.1* allele carrying maize W22 regulatory region and teosinte genic region (IN0949) is expressed about three-fold lower than a complete teosinte *ZmYAB2.1* allele (IN1207) [B]. Allele specific expression of *ZmYAB2.1* [C] is shown as log2 of teosinte allele expression level over maize allele expression level, and significance (p < 0.05) of each comparison (H_0_:μ=0) is indicated by an asterisk (*).

As a verification of this expression difference between the maize and teosinte alleles in the parental lines, we compared the expression of *ZmYAB2.1* in two RC-NILs. One RC-NIL (IN0949) carries a recombinant allele with the maize regulatory region and full-length maize:teosinte recombinant coding region without the dSpm insertion, while the other (IN1207) carries teosinte regulatory and coding regions (Supporting information, Figure S3). Interestingly, we observed expression of *ZmYAB2.1* in both of these two lines, but the expression is significantly higher with the teosinte allele (IN1207) as compared to the recombinant allele with the maize regulatory region (IN0949) (p = 0.00003) (Figure 2B). These results suggest that there is a difference in the 5’ *cis* regulatory elements of the maize (W22) and teosinte alleles. The maize *cis* element engenders lower expression than the teosinte *cis* element. The lack of *ZmYAB2.1* expression in T1L_D17 with the W22 allele presented above is likely a result of the dSpm insertion. As shown in Figure 1, T1L_D17 and IN0949 both show the maize class of ear internode length, while T1L_C07 and IN1207 both show the teosinte class of internode length. These results suggest that high expression of *ZmYAB2.1* confers longer ear internode length, while low or null expression of *ZmYAB2.1* confers shorter ear internode length.

In addition, we performed another expression analysis using an allele specific expression assay in plants heterozygous for different combinations of four maize (A619, Oh43, MR12, MR19) and two teosinte (TIL01, TIL11) alleles of *ZmYAB2.1.* We calculated the allele specific expression by taking log2 of the ratio of teosinte allele expression level over maize allele expression level. We observed higher expression of the teosinte *ZmYAB2.1* allele over the maize *ZmYAB2.1* allele in four of the six F1 hybrids with different allelic combinations (Figure 2C). One F_1_ showed higher expression of the teosinte allele, although the difference was not statistically significantly different from equivalent expression. Another F_1_ showed fully equivalent expression of the maize and teosinte alleles (Figure 2C). These results suggest that teosinte alleles are generally more highly expressed than maize alleles, however they also show some evidence for functional diversity among alleles within both maize and teosinte.

### Interactions with tb1

A previous study suggested that *tb1* is epistatic to *etb1.2* based on phenotypic data alone (Studer and Doebley 2011). We attempted to verify the epistasis by comparing the expression levels of *ZmYAB2.1* and *tb1* to ear internode lengths with respect to the genotypes at *ZmYAB2.1* and *tb1.* We defined the four different classes of genotypes at *ZmYAB2.1* and *tb1* as MM, TM, MT, and TT where the first letter represents maize (M) or teosinte (T) genotype at *ZmYAB2.1* and the second letter represents maize (M) or teosinte (T) genotype at *tb1.* Among these four classes, we observed a significant interaction effect between *ZmYAB2.1* and *tb1* for both ear internode length (p = 0.000107) and *ZmYAB2.1* expression (p = 0.003090) (Figure 3). For *ZmYAB2.1* expression, MM and MT classes show no detectable expression as expected, while TT class show significantly higher expression than TM class (p = 0.0152) (Figure 3A). This result indicates that the genotype at *tb1* has an effect on *ZmYAB2.1* expression. For *tb1* expression, MM and TM classes show similar expression level, and MT and TT classes show similar expression level albeit significantly lower than MM and TM classes (p = 0.00015) (Figure 3B). These observations indicate that *ZmYAB2.1* does not affect *tb1* expression.

**Figure 3.**
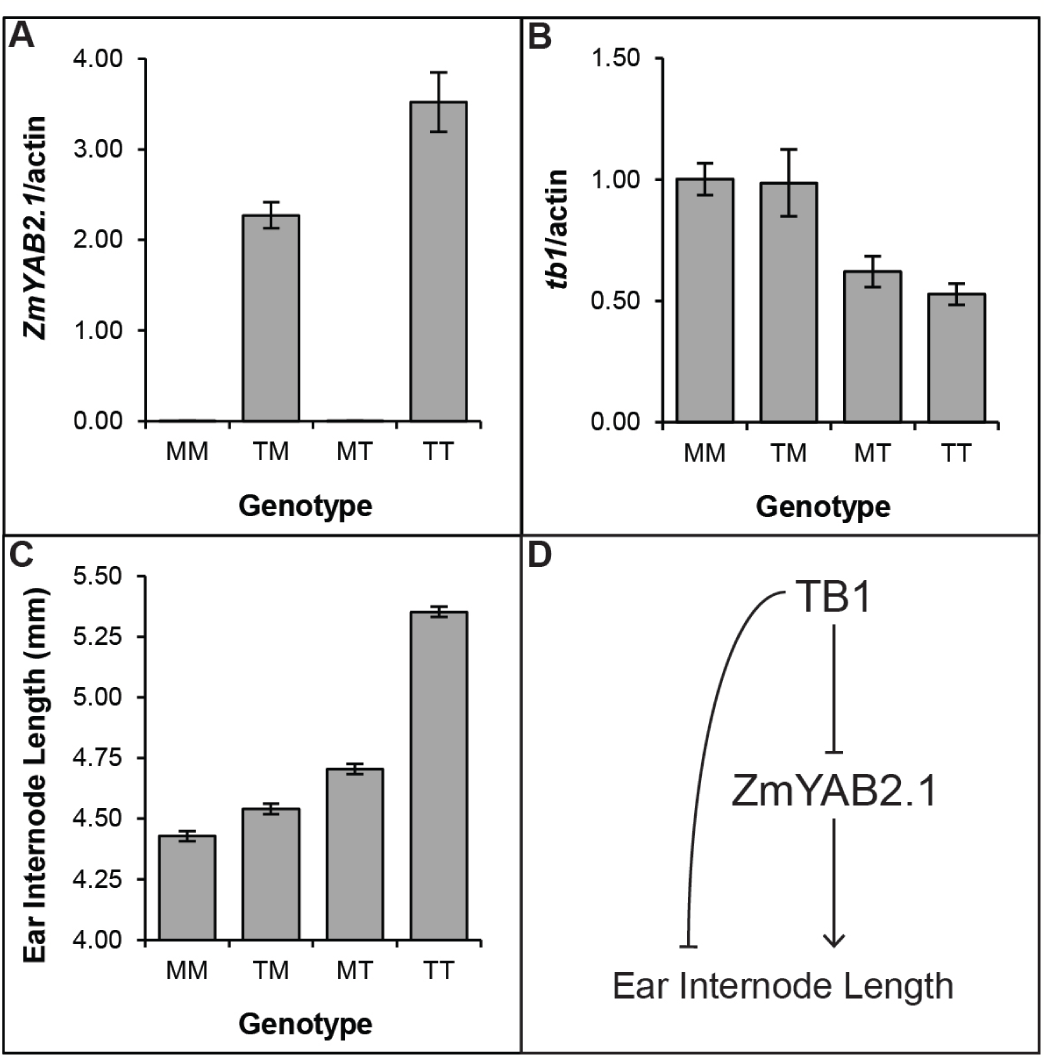
Expression levels of *ZmYAB2.1* [A] and *tb1* [B], and ear internode lengths [C] of four different genotype classes. Genotype of *ZmYAB2.1* is denoted on the left while genotype of *tb1* is denoted on the right (e.g. MT = maize allele at *ZmYAB2.1,* teosinte allele at *tb1).* Expression level of *ZmYAB2.1* is affected by *tb1* genotype [A]; however, expression level of *tb1* is unaffected by *ZmYAB2.1* genotype [B]. Pearson’s correlation is 0.72 between *ZmYAB2.1* expression level and ear internode length, and -0.69 between *tb1* expression level and ear internode length. Based on the expression and phenotype data, the most plausible model [D] involves 2 independent pathways. The first pathway begins with TB1 as a negative regulator of ZmYAB2.1, while ZmYAB2.1 is a positive regulator of ear internode length (as seen with the effect of *tb1* expression level on *ZmYAB2.1* expression level). The second pathway occurs independent of ZmYAB2.1 as TB1 negatively regulates ear internode length (as seen with the effect of *tb1* expression level on ear internode length when *ZmYAB2.1* expression is null). All interactions suggested here could be direct or indirect.

We also tested for significant interaction effects (epistasis) on ear internode length. Here, each of the four genotypic classes is significantly different from the others, with increasing ear internode length starting from MM, TM, MT, to TT classes (Figure 3C). However, the difference between MM and TM is small as compared to the difference between MT and TT which is much larger. Using the linear model as described in Materials and Methods, we identified a significant interaction term between *ZmYAB2.1* and *tb1* on ear internode length (p = 0.000107). The genotypic means for the four classes and the significant interaction term suggest that the genotype at *tb1* (M or T) has a larger effect on trait value when the teosinte allele is present at *ZmYAB2.1,* but a much smaller effect when the maize allele is present at *ZmYAB2.1* (Figure 3C). Based on these results, we see that the interaction between the teosinte *ZmYAB2.1* allele and teosinte *tb1* allele leads to higher *ZmYAB2.1* expression, and subsequently longer ear internode length.

The combination of expression and phenotypic data allow us to propose the following model. Expression level of *ZmYAB2.1* is dependent upon both the *ZmYAB2.1* and *tb1* genotypes, while expression level of tb1 is only dependent upon its own genotype. Our observations suggests that TB1 acts as a repressor of *ZmYAB2.1* directly or indirectly (Figure 3D). We thus propose that TB1 negatively regulates *ZmYAB2.1* expression, while ZmYAB2.1 positively regulates genes responsible for ear internode elongation (Figure 3D). However, this model is not complete as the difference in ear internode length between the MM and MT classes with null *ZmYAB2.1* expression is unexplained. This result suggests that an additional pathway through which TB1 independently regulates other genes responsible for ear internode elongation (Figure 3D).

### in situ Hybridization

In order to observe the spatial expression domain of *ZmYAB2.1* and any potential difference between the maize and teosinte alleles of *ZmYAB2.1,* we performed *in situ* hybridization on ears of both maize and teosinte alleles in both maize and teosinte genetic background (Figure 4). In our initial hybridizations, we used a mix of antisense probes containing the 3’ UTR probe, and either a coding sequence probe or a 5’ UTR probe, under standard hybridization conditions. In maize ears containing the maize allele (T1L_D17) (Figure 4A) as well as the teosinte allele (T1L_C07) (Figure 4B), we observed expression in a narrow band of cells subtending the spikelet pair. Similar results were seen in teosinte ears with the maize allele (TIL09) (Figure 4C) and the teosinte allele (TIL11) (Figure 4D), where expression was clearly localized to the future abscission zone between individual fruit-cases. Maize ears lack an abscission zone and considering the dramatic morphological changes resulting from domestication it can be difficult to identify homologous regions in the maize compared to teosinte ears. However, given the similar expression in a narrow band of cells in both maize and teosinte, we reason that the region of expression in maize is homologous to the abscission zone, indicating that a vestigial abscission zone is present in the maize ear.

**Figure 4.**
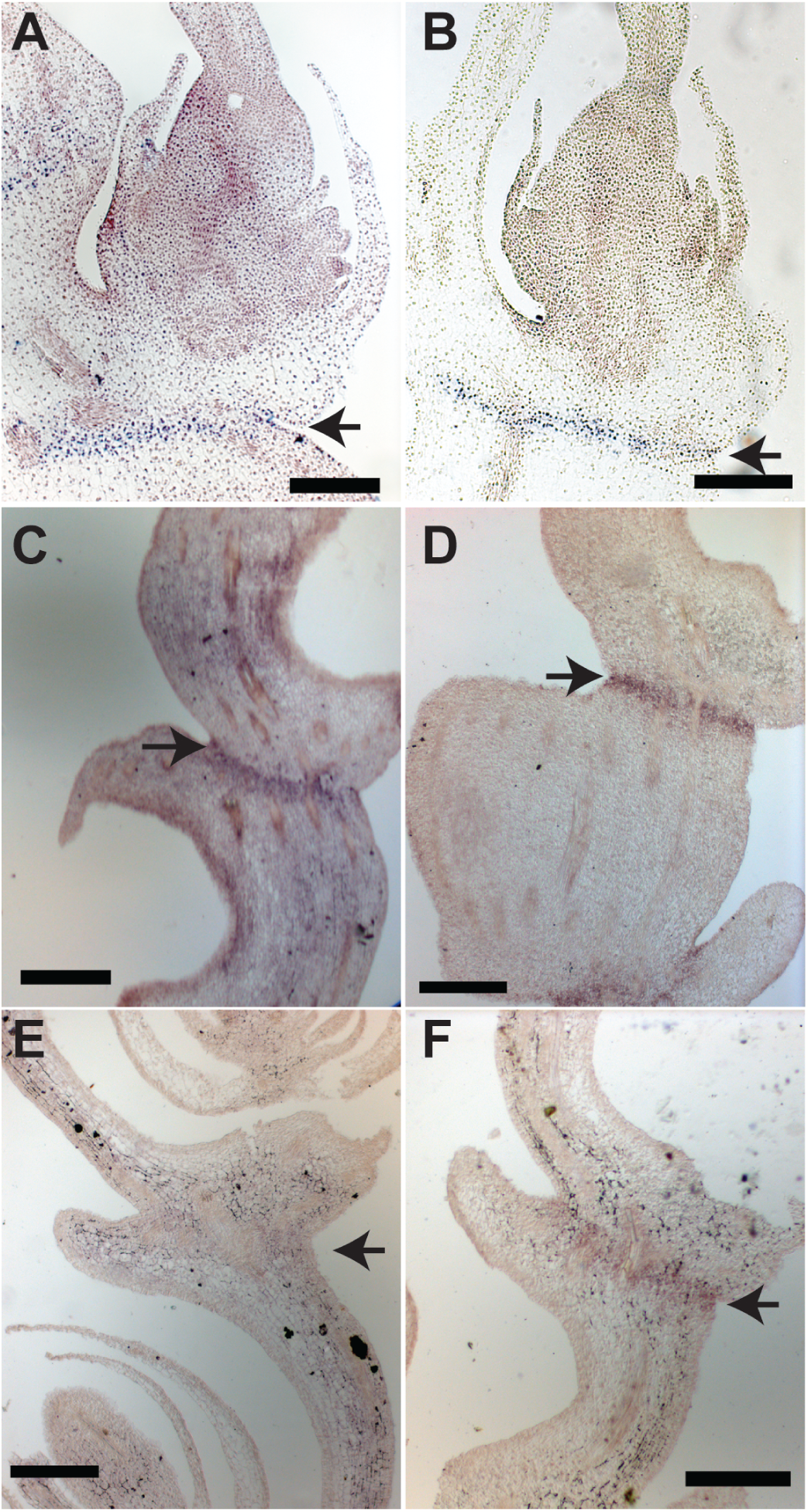
*In situ* RNA hybridization of *ZmYAB2.1* in maize and teosinte. A-D were hybridized with an anti-sense probe including both the coding sequence and 3’UTR. [A] is a maize ear with the maize allele of *ZmYAB2.1* while [B] is a maize ear with the teosinte allele. Expression is observed in a narrow band subtending the spikelet pair. This same probe hybridized to teosinte ears with the maize allele [C] or the teosinte allele [D] also show a narrow band of expression in the future dehiscence zone. A probe specific to the 3’ UTR hybridized to teosinte ears with the maize allele [E] or the teosinte allele [F] only showed weak expression in the dehiscence zone with the teosinte allele. Arrows indicate dehiscence zone in teosinte ears [C-F] and the homologous region in maize [A-B]. The scale bars in all panels are 200*μ*m.

Considering that the qPCR results clearly show a lack of expression of the 3’ region of *ZmYAB2.1* maize allele, we were surprised that our probe hybridized similarly to both alleles. One possible explanation for this discrepancy between the qPCR as compared to *in situ* expression results is that a closely related paralog (GRMZM2G074124) might cross-hybridize to our probe. A comparison of the coding sequence of the paralogs shows that they are 87% identical, while their 3’UTR region is only 67 % identical. Thus, we repeated the hybridizations with a probe specific to the 3’ UTR while also raising the hybridization temperature to provide more specificity (Figure 4E–F; Supporting information, Figure S5). In teosinte ears, only the teosinte allele showed some weak expression in the abscission zone (Figure 4F), while the maize allele showed no evidence of expression (Figure 4E). Similar results were seen in maize ears (Supporting information, Figure S5). These results are consistent with the view that the teosinte allele is expressed in the abscission zone, while the dSpm insertion present in the maize allele abolishes expression of the 3’ end of the transcript. Given that the internode length was the trait mapped to *ZmYAB2.1,* we infer that expression in the abscission layer promotes internode elongation. In this regard, Tanaka *et al.* (2012) have suggested that the rice YABBY gene, *TONGARI-BOUSHI,* acts non-cell autonomously.

## Discussion

### YABBY Gene Function and Ear Internode Length

Previous studies in *Arabidopsis* revealed that the YABBY gene family act redundantly to regulate complex patterning and growth decisions in lateral organ development (Sawa *et al.* 1999; Bowman and Smyth 1999; Eshed *et al.* 1999; Villanueva *et al.* 1999; Bowman 2000; Sarojam *et al.* 2010; Finet *et al.* 2016). The YABBY gene family consists of five clades, namely CRABS CLAW (CRC), INNER NO OUTER (INO), YABBY3/FILAMENTOUS FLOWER (YAB3/FIL), YABBY2 (YAB2), and YABBY5 (YAB5) (Yamada *et al.* 2011). *FIL* and *YAB3* act together to promote leaf growth and expansion including some aspects of adaxial-abaxial polarity (Siegfried *et al.* 1999). While *YAB2* and *YAB5* have no apparent loss of function phenotype in *Arabidopsis*, both genes significantly enhance the leaf and meristem *fil* and *yab3* phenotypes (Sarojam *et al.* 2010), indicating that *YAB2* has a redundant role in early leaf growth and patterning. Although limited to floral development, both *INO* (Villanueva *et al.* 1999) and *CRC* (Alvarez and Smyth 1999; Bowman and Smyth 1999; Eshed *et al.* 1999) also directly regulate organ growth and expansion. While *YAB2* gene function has not been as clearly elucidated, it likely contributes to tissue growth and expansion roles common to all YABBY family members. Interestingly, the tissue growth functions of YABBY have been shown to correlate with TCP gene expression (Sarojam *et al.* 2010). As *tb1* is a TCP family member, this could help explain the apparent epistatic interaction between *ZmYAB2.1* and *tb1.*

Studies on *YAB2* homologs in other species including rice, sorghum and maize suggest that *YAB2* plays a bigger role in floral organ regulation than just lateral organ polarity specification (Jang *et al.* 2004; Toriba *et al.* 2007; Lin *et al.* 2012). Examples of these *YAB2* homologs are *OsYAB2* (previously known as *OsYABBYl* and *OsShl*) in rice, *Shatteringl (Shl*) in sorghum, and *ZmYAB2.5.1* (previously known as *ZmShl-5.l). OsYAB2, Shl* and *ZmYAB2.5.l* function in the maintenance of abscission layers in the inflorescence (Lin *et al.* 2012). In addition, molecular analyses of *OsYAB2* did not identify a role in polarity regulation of lateral organs (Toriba *et al.* 2007). Ectopic expression of *OsYAB2* led to an increase in stamen and carpel number, with no impact on abaxial-adaxial identity of lateral organs (Jang *et al.* 2004). Adding a role of ear internode length regulation to the functions for members of the YAB2 clade suggests a common theme for these homologs, that is, regulation of growth within flowers and inflorescences. Additionally, all the described *YAB2* homologs are actually domestication genes (Lin *et al.* 2012), signifying the importance of members of YAB2 clade in domestication.

### etb1.2 as a Genetic Background Factor

As suggested in the name, *enhancer of tb1.2, etb1.2* refers to a QTL that modifies the effects of *tb1* and contributes to a genetic background in which the effects of *tb1* are enhanced. Genes that contribute to genetic background are expected to interact epistatically with the focal gene (Wade 2002). Consistent with this expectation, our phenotypic and expression data (Figure 3) show that *ZmYAB2.1* and *tb1* function non-additively in the regulation of ear internode length and *ZmYAB2.1* expression. Using the MM class (homozygous maize at both *ZmYAB2.1* and *tb1*) as a reference point, the TT class has more than twice the effect on ear internode length than the MT and TM classes combined. Thus, we see an additive-by-additive epistatic effect (Crow and Kimura 1970) between *ZmYAB2.1* and *tb1.* Such an epistatic effect, along with more complicated additive-by-dominance and dominance-by-dominance epistatic effects, are the mechanisms underlying genetic background (Wade 2002). Consequently, *ZmYAB2.1* represents a component of the background that modulates the effects of *tb1,* and the identification of *ZmYAB2.1* is the first step in understanding the genetic background that is affecting *tb1*.

The QTL at *tb1* was first discovered as a large effect QTL that controls multiple traits related to both ear and plant architecture (Doebley *et al.* 1990; Doebley and Stec 1993). The observed effects include the regulation of internode length in both the ear and lateral branches, as well as a large effect on inflorescence sex, the partial conversion of the ear into a tassel. The effects of the *tb1* QTL are influenced by an epistatic interaction with a second QTL on chromosome 3 (Doebley *et al.* 1995). Notably, when the *tb1* QTL was introgressed alone into maize genetic background, the size of the effects on internode length and inflorescence sex were diminished in comparison with the joint introgression of the *tb1* QTL plus the additional QTL on chromosome 3. This prior demonstration of a QTL on chromosome 3 contributing to the genetic background for *tb1* and our characterization of *etb1.2,* highlight the importance of genetic background effects for *tb1.* Consistent with these observations, Vann *et al.* (2015) have recently shown that the maize allele of the *tb1* regulatory region can segregate in natural teosinte populations without a detectable effect on tillering which may indicate genetic background effects.

Studer and Doebley (2011) further investigated the *tb1* QTL by asking whether it would fractionate into multiple liked QTL. In the 63 cM region surrounding *tb1*, these authors detected four additional QTL affecting ear and inflorescence architecture traits. *etb1.2* is one of these four additional QTL, and in addition to being closely linked to *tb1, etb1.2* is a background gene that modifies the effect of *tb1* through an epistatic interaction. The close linkage of *tb1* and *etb1.2* might have been a factor favoring selection on these two genes during domestication. Le Thierry D’Ennequin *et al.* (1999) have proposed that selection during domestication can favor tightly linked genes in outcrossing species when there is gene flow between the incipient crop and its wild ancestor, conditions that apply to the maize-teosinte domestication system.

### Nature of Selection at ZmYAB1.2

The signal of selection at *ZmYAB1.2* is much weaker than that observed with other maize domestication genes. Our fine-mapping places the causative site between the first exon and 68 kb upstream. To assay this region for selection evidence, we tested for selection at the distal (-68 kb), proximal (-34 kb) and 5’ region of *ZmYAB2.1.* We observed selection evidence only in the form of a single significant HKA test at the 5’ region (Table 1), which includes 5’ UTR and Exon 1. These results are very different from those with two other maize domestication genes (*tb1* and *tga1),* both of which show strong selection evidence near their respective causative lesions (Wang *et al.* 2005; Studer *et al.* 2011). The weak signature of selection might indicate either (1) the single positive HKA test is a false positive, (2) the 5’ region is too distant (relative to the local recombination rate) from the selection target site to detect a strong signal, or (3) *ZmYAB2.1* was the target of a soft sweep at or near its 5’ region (Innan and Kim 2004; Hermisson and Pennings 2005). In a whole genome selection scan, Hufford *et al.* (2012) did not observe selection evidence at *ZmYAB2.1,* suggesting either no selection or selection too weak to leave a signature. Overall, the evidence for selection at *ZmYAB2.1* must be considered weak at best.

Assuming that the significant HKA test for 5’ region is not a false positive, the likely target of selection would be an upstream *cis* regulatory element (CRE). Exon 1 lacks non-synonymous differences between our maize and teosinte alleles, so Exon 1 seems unlikely to harbor the causative polymorphism. The significant HKA test for 5’ region would then reflect selection on a linked CRE. We see no evidence for selection at the -34 kb proximal region, thus the proposed CRE should lie between this proximal region and Exon 1. The lack of selection evidence at the proximal region can be explained because this region shows very little linkage disequilibrium with the 5’ region (Supporting information, Figure S9) and thus selection on a CRE close to 5’ region is not apt to have left a signature of selection at the proximal region. We were unable to develop any PCR primer sets to assay sites between the first exon and the -34 kb proximal region.

Although the evidence for selection on a CRE upstream of *ZmYAB2.1* is weak, this evidence is augmented by the observation that a sample of multiple teosinte alleles of *ZmYAB2.1* are expressed more highly than a sample of multiple maize alleles, consistent with the hypothesis that selection acted on a CRE. Our expression data for three teosinte and five maize lines, show the teosinte allele to be expressed higher than the maize allele in all comparisons except Oh43 × TIL01 for which case equivalent expression was observed (Figure 2). Lemmon *et al.* (2014) reported allele specific expression at *ZmYAB2.1* for three maize and three teosinte lines, and their data also show that the teosinte alleles are expressed at a higher level than the maize allele (Supporting information, Table S7). Overall, these data suggest that maize was under selection for reduced expression of *ZmYAB2.1.*

If selection was acting on a CRE to reduce *ZmYAB2.1* expression, this may explain why four distinct loss of function alleles (LOF) are present in maize, some at moderately high frequencies (Table 2). Each of these LOF alleles occurs at low frequency in teosinte (~0-2 %) but higher frequency in maize landraces or inbreds (~4-22 %). The LOF alleles are geographically distributed in maize outside the initial area of maize domestication in southwest Mexico (van Heerwaarden *et al.* 2011), suggesting that they arose post-domestication (Supporting information, Figure S2). Their presence at low frequency in teosinte can be explained by introgression from maize into teosinte. The presence of a mix of low-expression and LOF alleles in maize suggests that loss/reduction of *ZmYAB2.1* function is beneficial in maize. Reduced or lost function would produce shorter internodes in the ear and more tightly packed kernels. Together the selection tests, expression data and LOF alleles can be explained by a model that a CRE at *ZmYAB2.1* was initially selected for lower expression, and then as maize spread out of it center of origin, multiple independent LOF alleles accumulated in different geographic regions.

The presence of multiple LOF alleles at *ZmYAB2.1* is reminiscent of allelic diversity at some other domestication-related genes. In maize, five different LOF alleles of the *sugary1 (su1*) gene have been identified to confer sweetness in sweet maize varieties (Tracy *et al.* 2006). Similar to *ZmYAB2.1* alleles (Supporting information, Figure S2), these five different *su1* alleles are also widely distributed based on geographical origin of each allele (Tracy *et al.* 2006). In addition, three different LOF alleles have been observed at *six-rowed spike 1 (Vrs1*) in barley that converts two-rowed barley into six-rowed barley (Komatsuda *et al.* 2007). Multiple LOF alleles of *waxy (wx*) have been identified to confer variation in amylose content in rice, resulting in glutinous and non-glutinous varieties of rice (Chen *et al.* 2014). Multiple LOF alleles of *5-O-glucosyltransferase* have been found to regulate anthocyanin contents in grapes (Yang *et al.* 2014). These examples stand in contrast domestication genes like *tb1* (Studer *et al.* 2011) and *tga1* (Wang *et al.* 2005) which have single domestication alleles that provide a gain of function.

### Pleiotropy at ZmYAB2.1

Aside from ear internode length regulation, *ZmYAB2.1* has also been previously suggested to regulate ear shattering (Lin *et al.* 2012). Such pleiotropic effect at *ZmYAB2.1* is not surprising given that ear internode elongation and shattering are biologically related traits (Khan *et al.* 2012). The shattering trait is interpreted as separation of two organs at an abscission zone, and it is an important mode of seed dispersal in wild species (Roeder and Yanofsky 2005). In *Arabidopsis, BREVIPEDICELLUS (BP), KNOTTED-LIKE FROM ARABIDOPSIS THALIANA 1 (KNAT1), KNAT2* and *KNAT6* are genes known to be involved in both inflorescence internode elongation and abscission zone formation (Smith and Hake 2003; Ragni *et al.* 2008; Shi *et al.*2011; Khan *et al.* 2012). Typically, abscission zone formation is restricted until seed maturity. However, upon seed maturity, both cell size and number increase at the abscission zone along with lignin deposition (Shi *et al.* 2011), resulting in the shattering. The increase in cell size and number would lead to internode elongation.

Despite *ZmYAB2.1* being a pleiotropic gene, we did not identify any significant variation for shattering trait in all the RC-NILs (data not shown). This result can be due to several reasons. Given that *ZmYAB2.1* is a gene of small effect, phenotyping accuracy can be crucial to tease the small difference apart. Unfortunately, our simple drop test method for scoring shattering trait is not as accurate as our method for scoring ear internode length. In addition, as discussed previously, genetic background plays an important role for characterizing a gene and our fine-mapping population lacks the correct genetic background for the shattering trait to be expressed. Also, of the two shattering QTLs that were previously mapped in maize, the LOD score for the QTL on chromosome 1 containing *ZmYAB2.1* is about five times lower than the QTL on chromosome 5 (Lin *et al.* 2012), indicating that *ZmYAB2.1* has a much smaller effect of these two QTL.

### Overview

We have taken a phenotype-to-gene mapping approach to better understand the nature of a factor, *etb1.2,* that contributes to the genetic background affecting the phenotypic effects of *tb1*. Surprisingly, or perhaps not, the gene underlying the factor was revealed to be a DNA-binding transcriptional regulator, *ZmYAB2.1,* like *tb1* itself. Transcription factors appear to be the most common class of genes involved in the evolution of morphological trait during domestication (Doebley 2006). Another surprise was that *ZmYAB2.1* had been previously identified as a likely gene contributing to the change from shattering to non-shattering ears during maize domestication (Lin *et al.* 2012). In addition to *etb1.2*, there are additional known factors that interact epistatically with *tb1* to affect multiple domestication traits (Doebley *et al.* 1995; Studer and Doebley 2011). The apparent intersection of multiple genes working in concert to regulate a suite of traits altered during domestication hints that selection during domestication operated to modify the function of a complex developmental network. Identifying other players in this network will be key to understand how genes like *tb1* and the background in which they work have evolved in concert.

## Acknowledgement

We thank Jordan Michels, Zhihong Lang, Zachary Lemmon, Jesse Rucker, Elizabeth Buschert and Eric Rentmeester for helpful assistance throughout the entire project. This work was supported by the National Science Foundation, funding IOS1025869, IOS0820619, and IOS1238014.

